# Direct and Inverted Repeat stimulated excision (DIRex): Simple, single-step, and scar-free mutagenesis of bacterial genes

**DOI:** 10.1101/149534

**Authors:** Joakim Näsvall

## Abstract

The need for generating precisely designed mutations is common in genetics, biochemistry, and molecular biology. Here, I describe a new λ Red recombineering method (**D**irect and **I**nverted **R**epeat stimulated **ex**cision; DIRex) for fast and easy generation of single point mutations, small insertions or replacements as well as deletions of any size, in bacterial genes. The method does not leave any resistance marker or scar sequence and requires only one transformation to generate a semi-stable intermediate insertion mutant. Spontaneous excision of the intermediate efficiently and accurately generates the final mutant. In addition, the intermediate is transferable between strains by generalized transductions, enabling transfer of the mutation into multiple strains without repeating the recombineering step. Existing methods that can be used to accomplish similar results are either (i) more complicated to design, (ii) more limited in what mutation types can be made, or (iii) require expression of extrinsic factors in addition to λ Red. I demonstrate the utility of the method by generating several deletions, small insertions/replacements, and single nucleotide exchanges in *Escherichia coli* and *Salmonella enterica*. Furthermore, the design parameters that influence the excision frequency and the success rate of generating desired point mutations have been examined to determine design guidelines for optimal efficiency.

## Introduction

λ Red recombineering is a very powerful method for modifying bacterial chromosomes, plasmids, and BAC clones. Initially it was used to replace chromosomal genes by antibiotic resistance cassettes carried on PCR products [1,2]. While antibiotic resistance genes are very practical as selection markers it is often undesirable to leave selection markers behind. The limited number of antibiotic resistance markers can constrain the number of mutations that can be combined into a single strain. Hence, strain constructions may require careful planning and additional constructions to avoid running out of markers. One solution to this problem is the introduction of markers flanked by directly repeated Flp recombinase target (FRT) sites [2]. These markers can be excised by expressing the site-specific recombinase Flp, leaving a single copy of FRT behind (an FRT “scar”). A downside of this method is however that it creates the risk of making unintended genetic rearrangements due to recombination between FRT scars at different loci [2,3]. Another problem with this and previous methods is that both an FRT scar and antibiotic resistance cassettes can influence the expression of surrounding genes, resulting in unintended phenotypes and erroneous genotype to phenotype correlations. For example, we have observed a significant reduction in growth rate when an FRT scar was present at the end of the *Salmonella enterica rpsT* transcript, outside of the coding sequence [4]. To account for such unintentional effects that are difficult to predict and infer the true effect of a particular mutation, it is standard practice to always compare isogenic strains (i.e. strains that differ by one single mutation) in all experiments [5]. But the more mutations that are added to an experiment, the more strains are needed as isogenic controls in order to account for all markers. This may limit the number of mutations that can be included in multiplex experiments and to avoid these problems, it is of great importance to find methods that do not leave selection markers or scar sequences behind in the final strains.

Several recombineering methods exist for generation marker-free and scar-free mutants. These existing strategies either require (i) screening in the absence of a selectable phenotype, or (ii) overlap-extension PCR to generate a complex locus-specific cassette, or (iii) expression of additional factors to induce chromosome breaks (e.g. I-SceI or CRISPR/Cas9 plus a guide RNA), or (iv) several transformation steps [6–14].

Some of the above shortcomings can be overcome by utilizing the inherent instability of inverted repeat (IR) sequences flanked by direct repeats (DR). In both *Escherichia coli* and *Saccharomyces cerevisiae* long inverted repeats are excised at high frequency by deletions between flanking direct repeats, leaving one copy of the DR sequence [15–17]. This has been exploited by yeast geneticists in a method termed Mutagenic Inverted Repeat Assisted Genome Engineering (MIRAGE) [18]. Tear et al. [19] demonstrated a λ Red based method very similar to MIRAGE in *E. coli* that require designed locus derived IRs to be constructed by overlap extension PCR (OE-PCR) for every mutant construction. Using locus derived IRs restricts the method to generation of relatively large deletions, as the inverted repeats by necessity need to be within the deleted area. This also efficiently prevents generation of point mutations and small insertions or replacements.

Here I expand this toolkit with a new method, termed **D**irect- and **I**nverted **R**epeat stimulated **ex**cision (DIRex). This method is based on the same basic principle as MIRAGE, but does not require any OE-PCR to generate the recombinogenic DNA for recombineering. Briefly, a selectable and counter selectable cassette containing long terminal IRs is placed between two short locus-derived or synthetic DRs, generating a semi-stable intermediate insertion (DIRex intermediate). DIRex intermediates are stable enough to be transferable with generalized transduction, but unstable enough that the designed mutants are easily isolated by subsequent counter-selection. With DIRex it is possible to generate deletions and point mutations, and even small insertions or replacements as long as the introduced sequences can be included in long PCR primers. The utility of DIRex is demonstrated through deletion of several genes, sequence replacements, and for introduction of single nucleotide substitutions. Observations made during method development may also have implications for the understanding of the mechanism of λ Red mediated dsDNA recombination.

## Results

### Overview of DIRex

The DIRex method, outlined in Fig 1 and Fig 2, involves a single λ Red recombineering step to generate a semi-stable DIRex intermediate followed by a quick and simple procedure to isolate the final mutant. Once the DIRex intermediate is constructed, it can optionally be transferred to other strains by generalized transduction (Fig 1B). The DIRex intermediate consists of a selectable and counter-selectable marker, flanked on both sides by long identical IRs (Fig 2C). Flanking these IRs are locus derived DRs that are short enough to be contained in the primers used for PCR. The DRs provide enough homology for spontaneous and precise excision of the semi-stable insertion. The DRs needed for excision can be designed to consist of locus derived sequences containing or flanking the desired mutation (Fig 3). If each of the “outer” primers contain 15 extra nucleotides between the 5’-recombinogenic homology extension and the 3’-template annealing portion, a 30 bp DR with the designed mutation in the middle can be constructed (Fig 3A).

**Fig 1.**
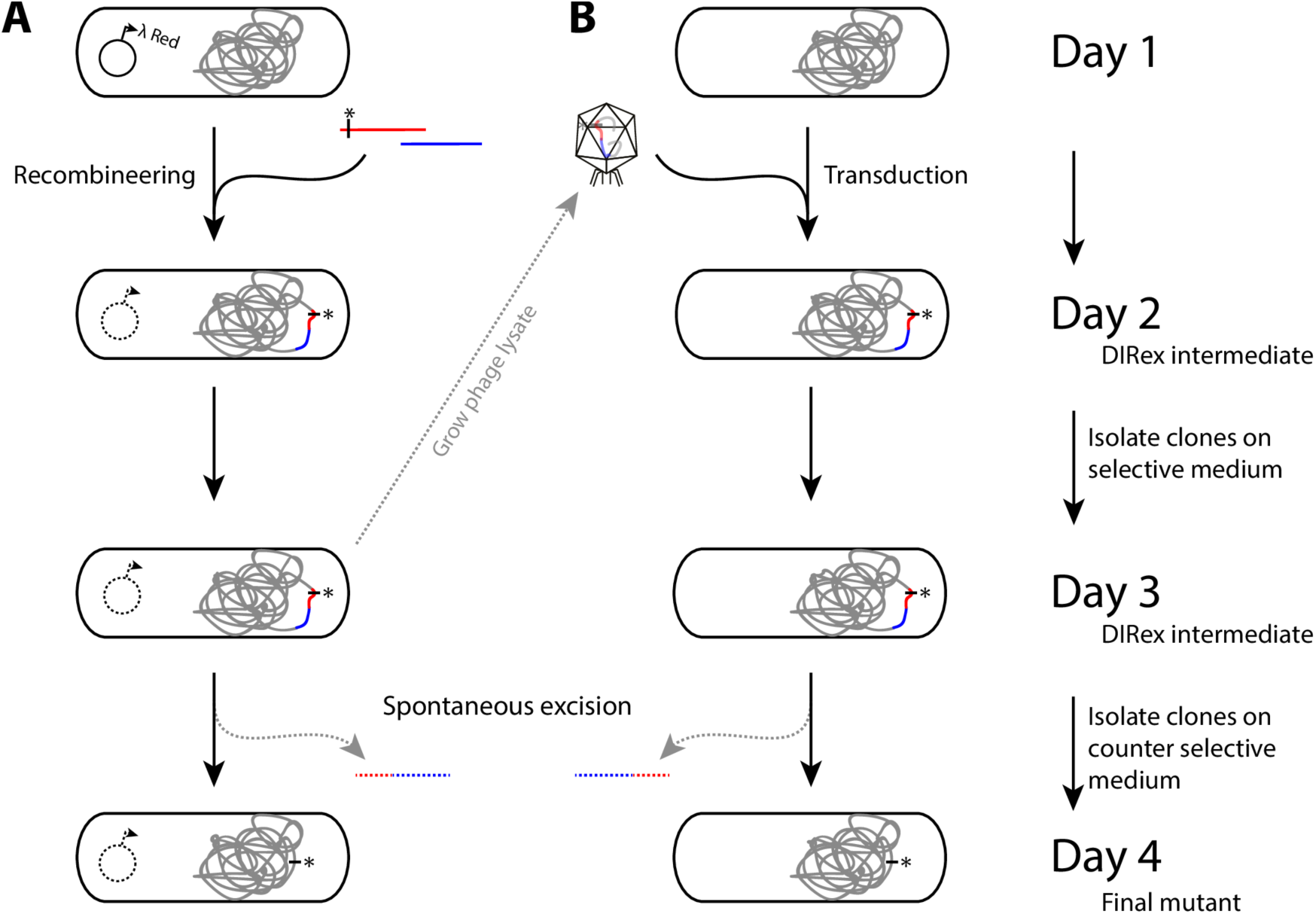
Outline of the method. (A) Generation of a designed mutation in three days using DIRex. (Day 1) A culture expressing λ Red is transformed with two PCR products to generate a semi-stable DIRex intermediate containing a selectable and counter selectable cassette. See Fig 2 for more details. (Day 2) Transformants are isolated and colony-purified on selective medium. (Day 3) Colonies growing on selective medium are picked and streaked on counter-selective medium. (Day 4) Nearly 100% of colonies growing on counter selective media contain the designed mutation. (B) Transferring a previously constructed mutation into another strain by generalized transduction. A phage lysate grown on a strain containing a DIRex intermediate is used as donor in the transduction. The steps involved are the same as in (A) except for a transduction instead of a recombineering step on day 1.

**Fig 2.**
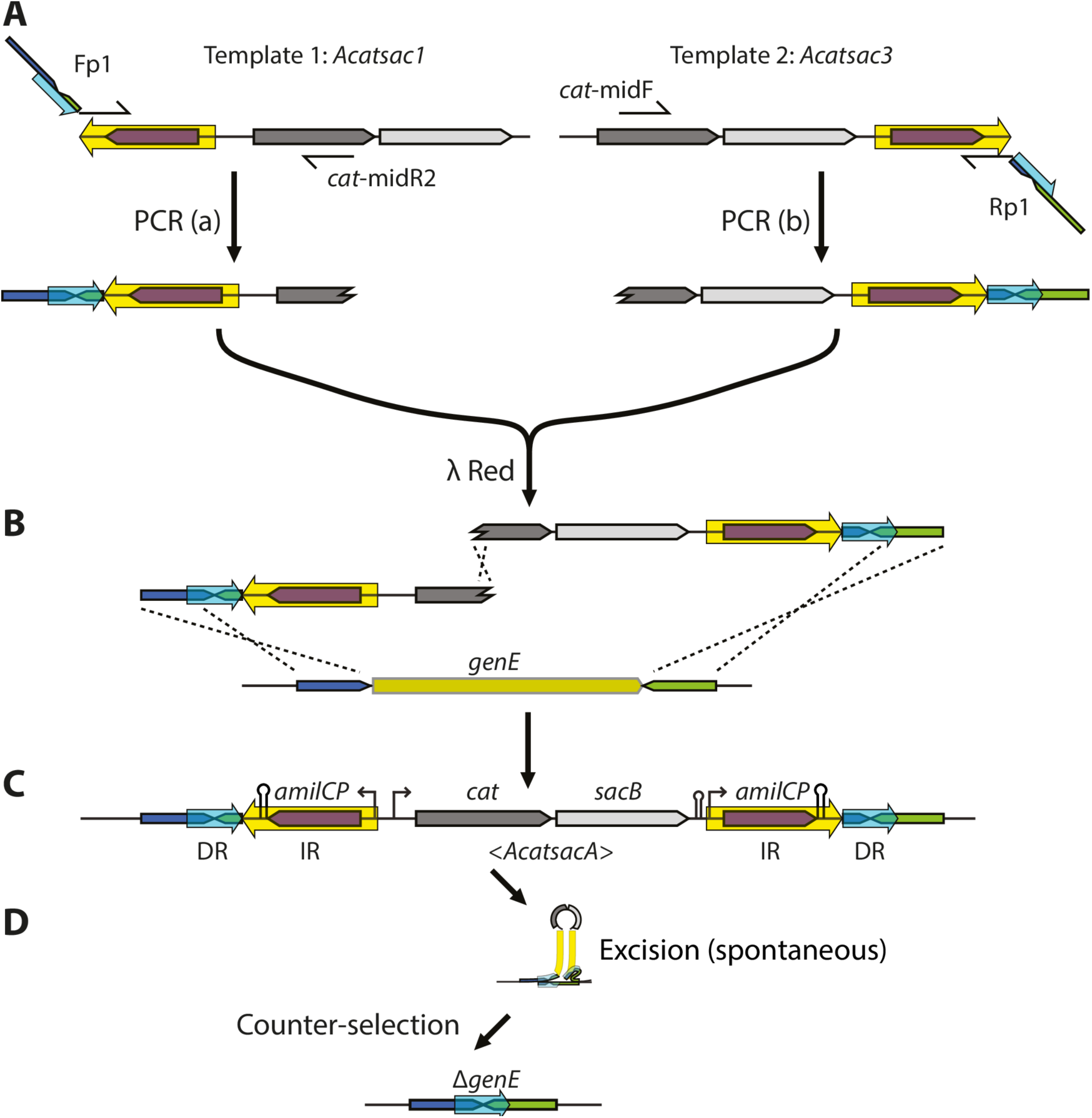
Overview of the method. The method is illustrated with an example for generating a precise deletion of a hypothetical gene. (A) Two overlapping “half-cassettes” are generated in separate PCR reactions (which can be run in parallel in the same PCR cycler) using one locus specific long primer “Fp1” or “Rp1” in combination with the cassette specific primers “*cat*-midR2” or “*cat*-midF”, respectively. Each PCR fragment contain one copy of the IR (yellow arrow) and DR (light blue arrrow), as well as one of the recombinogenic 5’-homology extensions. The templates (*Acatsac1* and *Acatsac3*) differ in the location and orientation of the IR sequence, which contains the gene encoding the blue chromoprotein AmilCP. (B) The two “half-cassettes” are mixed in equimolar amounts and electroporated into λ Red induced cells. For formation of a functional *cat* gene recombination has to occur between the recombinogenic ends and the chromosome, as well as in the sequence overlap between the two “half-cassettes”. (C) The structure of the semi-stable DIRex intermediate. (D) The structure of the final deletion after spontaneous excision of the DIRex intermediate (See S3 Fig for a possible mechanism of excision).

**Fig 3.**
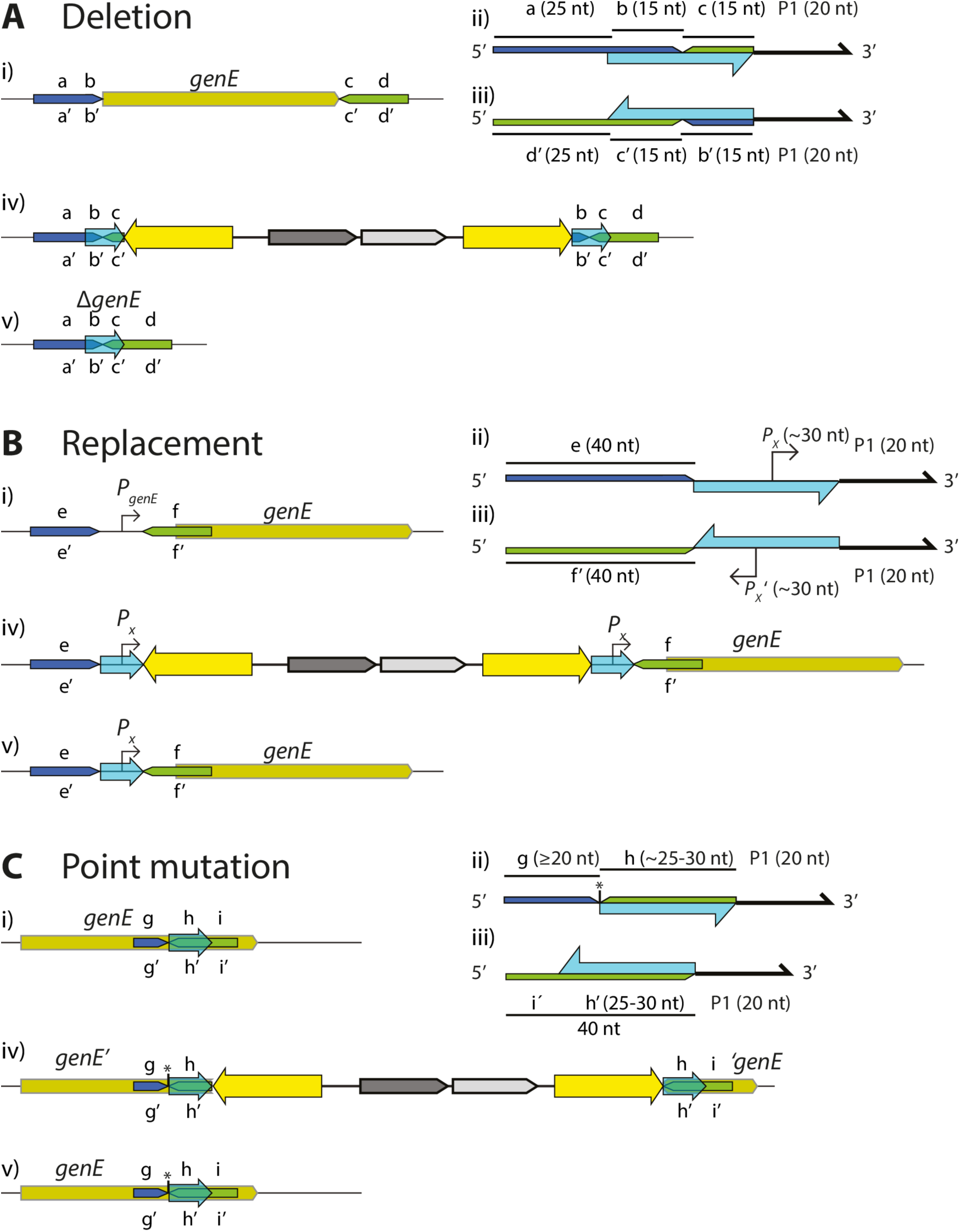
Primer design for DIRex. Light blue arrows indicate the location and orientation of DR sequences between which recombination is expected to occur. Sequence segments on the upper strand are labeled with lower case letters a – i, and on the lower strand the complementary sequences are labeled a’ – i’. (A) Using DIRex for deleting a gene. (A, i) Two 40 nt regions of recombineering homology (a-b, dark blue; c-d, green) are chosen on either side of the sequence to delete. (A, ii) The “left” oligo (upper strand) is designed with a 40 nt homology extension (composed of segment a - b) followed by 15 nts from the other side of the sequence to delete (segment c), and a 20 nt 3’ primer (P1). (A, iii) The “right” oligo (lower strand) is designed with a 40 nt homology extension (composed of segment d’ – c’) and 15 nts from the other side of the sequence to delete (segment b’), and a 20 nt 3’ primer (P1). (A, iv) The resulting DIRex intermediate, with two identical 30 nt DR sequences (b – c), each containing the designed deletion junction. (v) The designed deletion after excision of the DIRex intermediate. (B) Using DIRex for replacing a native sequence with a designed sequence (in this example replacing the promoter *P*_*geneE*_ with another promoter, *P*_*x*_). (B, i) Two 40 nt regions of recombineering homology (e, dark blue; f, green) are chosen on either side of the sequence to replace. (B, ii) The “left” oligo (upper strand) is designed with a 40 nt homology extension (composed of segment e) followed by the sequence to replace it with (in this example a ∼30 nt sequence containing a promoter, *P*_*x*_), and a 20 nt 3’ primer (P1). (B, iii) The “right” oligo (lower strand) is designed with a 40 nt homology extension (composed of segment f’) followed by the sequence to replace it with (the reverse complement of *P*_*x*_), and a 20 nt 3’ primer (P1). (B, iv) The resulting DIRex intermediate, with two identical 30 nt DR sequences, each composed of the replacing sequence (*P*_*x*_). (B, v) The designed replacement after excision of the DIRex intermediate. (C) Using DIRex for introducing a point mutation. The desired mutation is marked with an asterisk. (C, i) Two regions of recombineering homology (g-h and h-i) are chosen on either side of the point mutation. (C, ii) The “left” oligo (upper strand) is designed with a 45 – 50 nt homology extension (composed of segment g, the desired point, and segment h), and a 20 nt 3’ primer (P1). (C, iii) The “right” oligo (lower strand) is designed with a 40 nt homology extension (composed of segment i’ – h’), and a 20 nt 3’ primer (P1). The homology extension ends just next to the nucleotide(s) to be changed (C, iv) The resulting DIRex intermediate, with two identical 25 – 30 nt DR sequences, with the mutation next to the “left” DR sequence. (C, v) The designed mutant after excision of the DIRex intermediate. See S1 Fig for specific examples.

Our recent observation that recombineering with overlapping PCR fragments is as efficient as recombineering with a single, similar sized PCR product enables generation of complex insertions with repetitive sequences without the need for OE-PCR [20]. To generate a DIRex intermediate with inverted copies of an ∼840 bp DNA sequence at both chromosome-cassette junctions, two partial cassettes are generated in separate PCR reactions (Fig 2). One PCR product contains one of the recombinogenic extensions, one copy of the locus-derived DR, one copy of the IR plus a 3’ truncated *cat* gene (chloramphenicol resistance; Fig 2A, “PCR(a)”. The other PCR product contains a 5’ truncated *cat* gene, the *Bacillus subtilis sacB* gene (conferring sensitivity to sucrose), the second copy of the IR sequence, the second copy of the DR and the other recombinogenic extension (Fig 2A, “PCR(b)”). As the two truncated *cat* genes overlap by 277 bp recombination between the two PCR products regenerates a functional *cat* gene during the λ Red mediated recombination (Fig 2B). The homology extensions used for λ Red recombineering are included in the “outer” PCR primers, and contain sequences that are used to generate small (∼30 bp) DRs at the end of the large IR sequence. The IRs contain a copy of the gene encoding a blue chromoprotein (AmilCP from *Acropora millepora*), providing a simple colorimetric screen against most false positives. To generate these PCR products two templates and four primers are needed; two “outer” primers (Fig 2A, “Fp1” and “Rp1”) that need to be designed specifically for each construct, and which both contain the identical 3’ 20-mers as well as complementary sequences that generate the DRs that surround the inserted cassette, and two “inner” primers (Fig 2A, “*cat*-midR2” and “*cat*-midF”) that are specific to the antibiotic resistance gene and can be re-used for any construction. The resulting cassette in the DIRex intermediate (Fig 2C) is referred to as <*AcatsacA*> (abbreviated from *amilCP*-*cat*-*sacB*-*amilCP*, with the greater-than and smaller-than signs indicating the IRs). Excluding the time for primer design and synthesis, PCR verification and sequencing, clones containing the intended modification without any selectable marker or scar sequence can be constructed in as little as three days (transformation – cleanup – segregation; Fig 1).

### Direct repeats of 25 bp are enough for high frequency of accurate excision

Tear *et al.* achieved high accuracy of excision with 25 bp DRs, but did not provide any explanation as to why this size was chosen [19]. To examine if and how the DR size affects excision frequencies, DIRex intermediates flanked by 20, 25 or 30 bp DRs were generated in the *hisA* gene in *S. enterica* (Fig 4A). Several colony-purified blue, Cam^R^, His^-^ clones from each transformation were colony-purified on sucrose selection plates. All initially blue clones generated white colonies on sucrose plates, with some interspersed blue colonies. To estimate the frequency of excision versus false positives (blue Suc^R^ colonies) dilutions of overnight cultures were plated on LA and sucrose plates to calculate the frequencies of white and blue segregants (Fig 4B). The frequencies of Suc^R^ cells were higher with longer DRs, and the frequencies of blue Suc^R^ clones that still had part of the DIRex intermediate were relatively constant and always lower than the number of white Suc^R^ colonies. With 25 and 30 bp DRs, the vast majority (>98%) of the Suc^R^ clones were white, while with the smallest DR size 22% of the colonies were blue.

**Fig 4.**
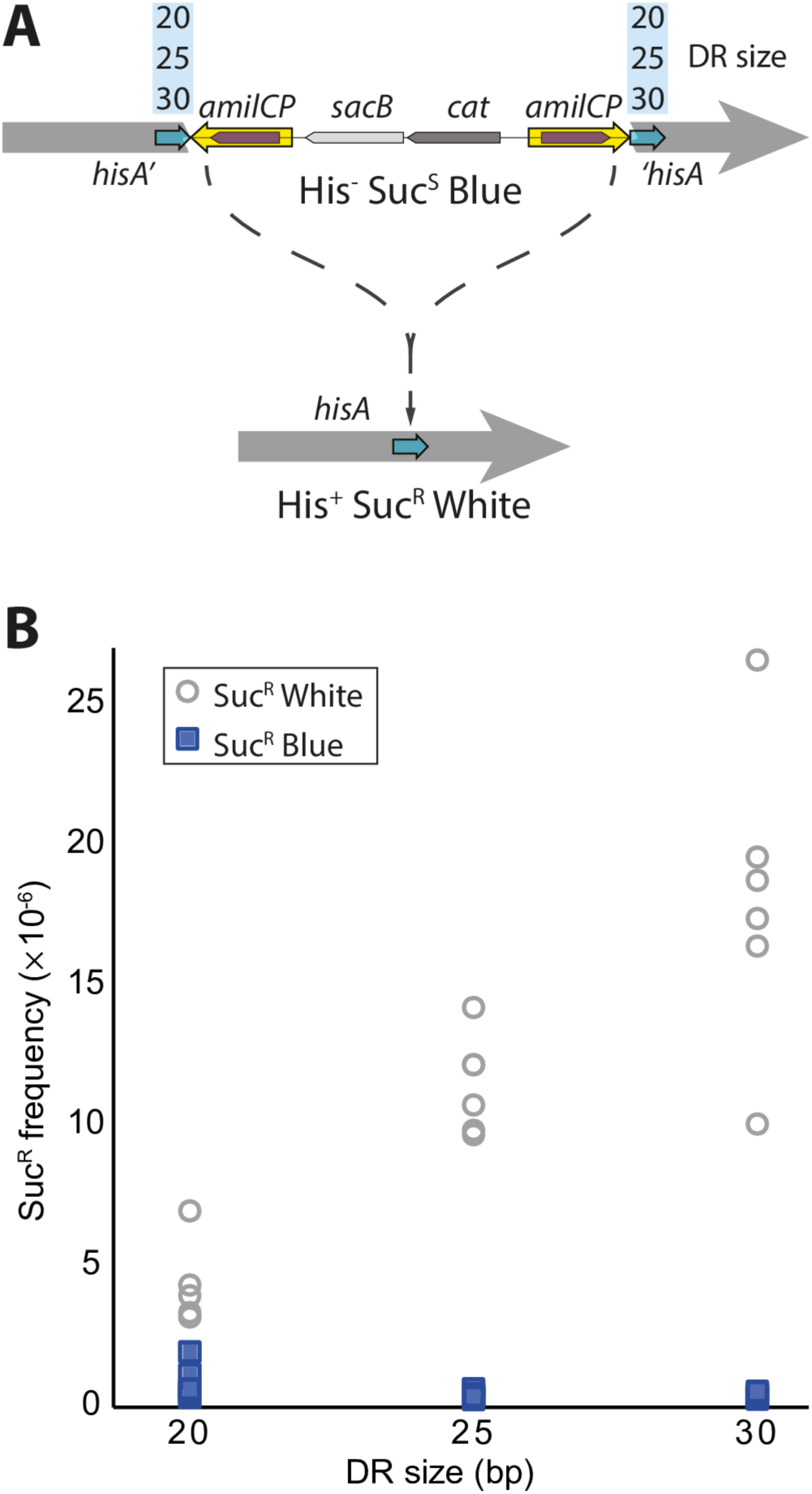
Excision frequencies increase with increasing DR size. (A) Assay for precise excision. The three DIRex constructs in *hisA* were identical except for having 20, 25 or 30 bp DRs (light blue arrows). When the DIRex intermediate is present the *hisA* gene is interrupted, and the cell is unable to grow in medium lacking histidine (His^-^). AmilCP confers blue color and SacB causes sensitivity to sucrose. Selection on sucrose selection plates allows only cells lacking a functional *sacB* gene to grow. If the cassette is excised a functional wild-type copy of *hisA* is restored and the cells lose the blue color and becomes sucrose resistant. For a more detailed view of the primers used in this experiment, see S1B Fig. (B) Frequencies of segregants (white) and false positives (blue) in six independent cultures of each of the three constructs.

As some of the false positives were expected to be deletions within the *cat*-*sacB* cassette that would produce white colonies, the smaller DR size could result in an unacceptably high frequency of false positive clones among the white Suc^R^ clones. To test this and verify that the majority of the white Suc^R^ clones were the result of precise excision of the DIRex intermediate, white Suc^R^ clones were screened for a functional *hisA* gene. One hundred white Suc^R^ colonies isolated from two separate cultures from each of the constructs were patched on M9 glucose plates. For all three constructs, at least 96% of the tested colonies were His^+^. As the frequencies of cells that had precisely excised the cassette was in the order of 10^−5^ per cfu, picking a single colony (∼10^8^ cells) and streaking for single colonies on a sucrose plate results in hundreds to thousands of segregants. However, although not found in this experiment, occasional “jackpots” with white false positive clones should be expected to occur.

### Using DIRex to make precise deletions or replacements

To delete a DNA sequence using DIRex, the two “outer” primers were designed so that the resulting insertion was flanked at both sides by 30 bp DRs containing the desired deletion junction (Fig 3A). Oligos for deleting several genes or genomic regions were designed (Fig 5 and S2 Table). In *S. enterica* the gene *ssrA* (tmRNA) (Fig 5A), the *gal* operon promoter region (Fig 5B) and the *araCBAD* genes (Fig 5C) were targeted. In *E. coli* a deletion of the *araFGH* operon (Fig 5D) was constructed. As one copy of the DR sequence remains after excision of the DIRex intermediate, generation of deletions and point mutations require the DR sequence to be locus derived. In other cases, it may be useful to leave an artificial sequence at the locus. As a demonstration of this, DIRex was used to replace the native (regulated) *araE* promoter (*P*_*araE*_) with a constitutive synthetic promoter (*P*_*J23106*_) in both *S. enterica* and *E. coli* (Fig 5E). For these constructions, primers were designed to introduce the *P*_*J23106*_ promoter as part of an artificial 40 bp DR, with 5’-recombinogenic extensions for deleting the native promoter. In addition, 933 bp of the 1017 bp *S. enterica galE* coding sequence was replaced with the 46 bp *lux* transcriptional terminator (*T*_*lux*_ [21]), using a similar design (Fig 5F). Similar designs could be used for inserting short sequences such as degradation tags, affinity purification tags, protein binding sites in DNA *etc*. For all of these constructions eight blue, Cam^R^ clones were colony purified in the presence of chloramphenicol, where after a single blue colony was streaked out on sucrose selection plates. In all cases this resulted in growth of white, Suc^R^ colonies and only a few false positive clones (blue, Suc^R^). Deletions and replacements were verified by PCR and sequencing. Most tested clones displayed the expected PCR fragment sizes and correct sequence with the exception of a few clones that contained additional mutations consistent with primer synthesis errors (single nucleotide deletions or insertions within the oligo-derived sequence; data not shown). To combine several mutations the Δ*araCBAD* Δ*P*_*araE*_::*J23106* and the Δ*araFGH* Δ*P*_*araE*_::*J23106* double mutants were constructed by P22 transductions in *S. enterica* and P1 transductions in *E. coli* respectively (S1 Table). To combine the very closely located Δ*galE*::*T*_*lux*_ and Δ*P*_*gal*_ mutations, Δ*P*_*gal*_::<*AcatsacA*> was transformed into a Δ*galE*::*T*_*lux*_ mutant (S1 Table).

**Fig 5.**
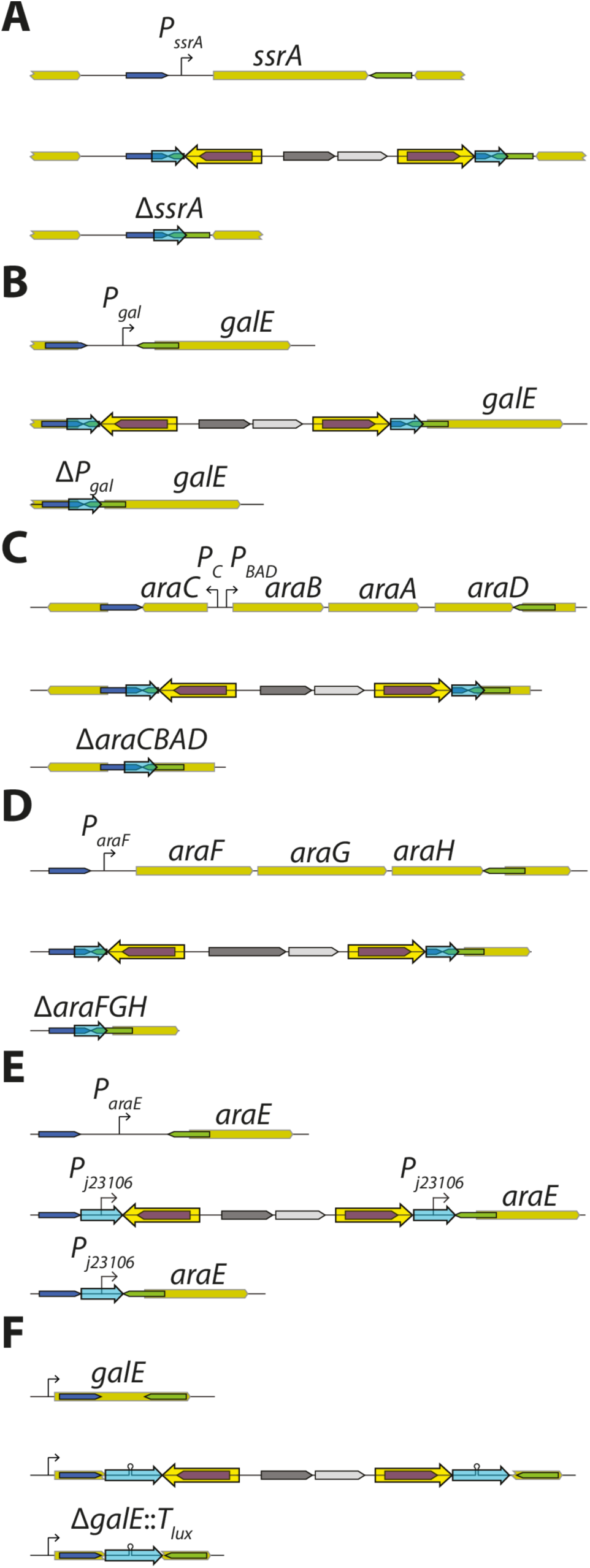
Use of DIRex to generate deletions and replacements. Dark blue and green areas indicate the recombinogenic homology arms and light blue arrows indicate the DRs used for excision. For each construct, the top line shows the wild-type arrangement, the middle line the DIRex intermediate and the bottom line the final mutant. (A) Deletion of the *S. enterica ssrA* gene, encoding tmRNA. (B) Deletion of the *S. enterica gal* operon promoter region. (C) Deletion of the *S. enterica araC*, *B*, *A* and *D* genes. (D) Deletion of the *E. coli araFGH* operon, encoding a high affinity L-arabinose transporter. (E) Replacement of the native *araE* promoter with a synthetic promoter (*P*_*J23106*_) in both *S. enterica* and *E. coli*. (F) Replacement of most of the *S. enterica galE* coding sequence with a *lux* transcriptional terminator.

### Using DIRex to generate point mutations

#### Most transformants segregate to only the designed allele

Several pairs of oligos were designed for correcting a mutation in the *hisA* gene. The mutation L169R was chosen since it allowed a simple screen for correct and incorrect transformants. The L169R allele of *hisA* has no detectable HisA activity but has a weak TrpF activity [22]. Hence, a strain lacking the native *trpF* gene and having the *hisA*(L169R) allele is capable of slow growth in the absence of tryptophan but not in the absence of histidine (phenotypically His^-^, Trp^+^). The *hisA*(L169R) mutants were transformed with DIRex constructs where the DRs contained sequences that would revert the mutation back to wild-type (a leucine codon; Fig 6A and S1A Fig) and restore histidine prototrophy. After loss of the inserted cassette by selection for sucrose resistance, correct transformants (*hisA*^+^, Leu 169) would have restored wild-type HisA function, but lost the weak TrpF activity (His^+^, Trp^-^). The strategy outlined in Fig 6B and S1A Fig was used to test the efficiency of generating the correct alleles when the mutation was present in the middle of both copies of the DRs. In this test 73% of the transformants segregated to only the intended allele, and most of the remaining transformants segregated to both possible alleles. Evidently, some transformants had the wanted allele (Leu 169) in only one of the DRs, while the other contained the parental allele (Arg 169). These transformants later segregated to generate both alleles, acting like heterozygotes. This suggests recombination during Red-mediated recombineering does not always result in introduction of the entire homology arm from the transforming fragment, leading to retention of the parental allele in a fraction of the transformants.

**Fig 6.**
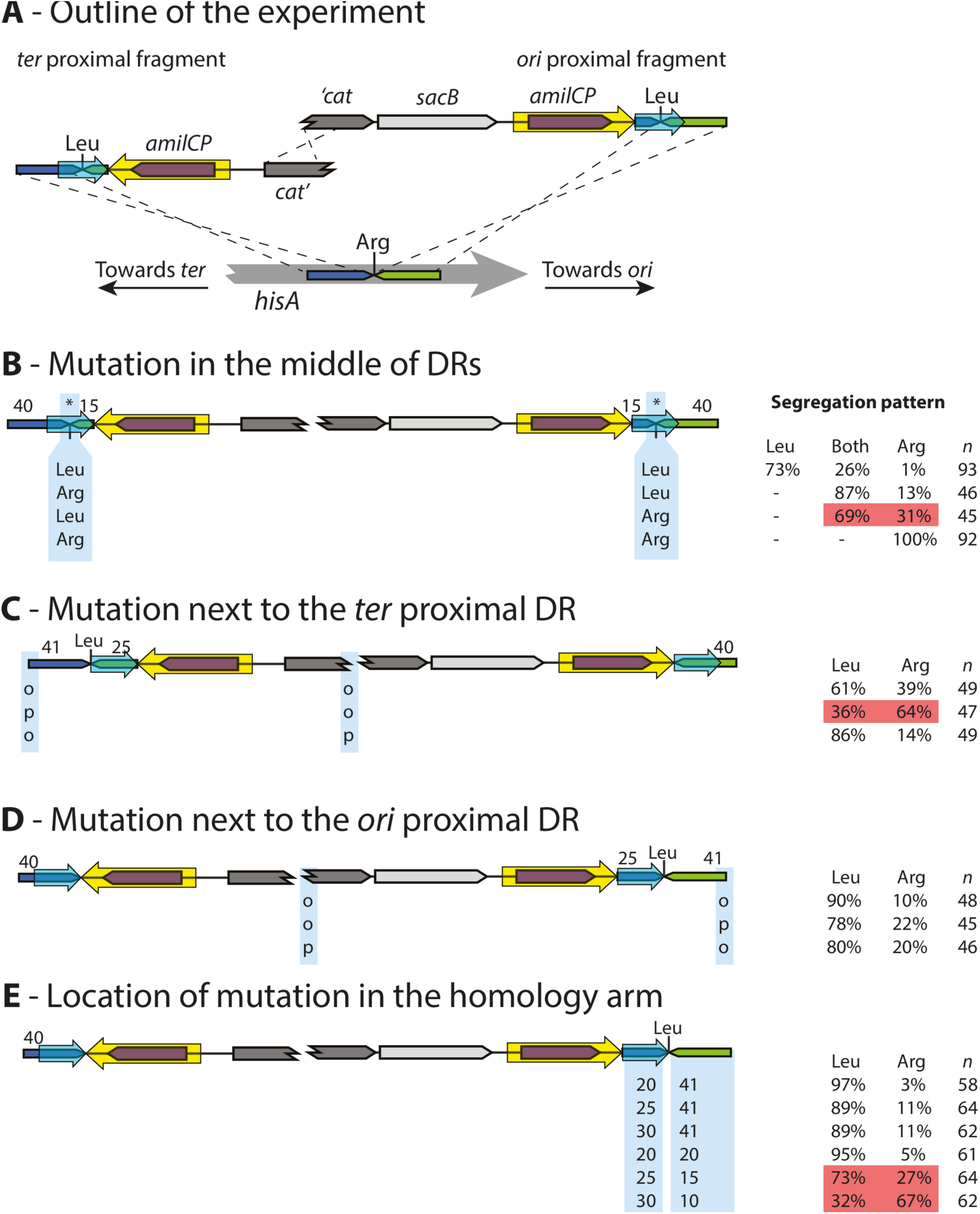
Alternative strategies for generating point mutations with DIRex. (A) Schematic view of the recombineering used for these tests. The recipient strain carries a L169R mutation in *hisA*, rendering it His^−^ (but Trp^+^). The incoming PCR fragments are designed to repair this mutation, making the wanted transformants (after segregation of the cassette) His^+^ (but Trp^−^). (B) Constructs with the mutation in the middle of the DRs. The site of the mutation is indicated in both DRs with asterisks. “Leu” and “Arg” below the DRs indicate which allele was present in the corresponding PCR product. In the table to the right, “Leu” indicates the fraction of transformants that only segregated to the L169 (wanted) allele, “Both” the fraction of transformants that segregated to both alleles, and “Arg” the fraction of transformants that only segregated to the R169 (parental) allele. (C) Constructs with the mutation next to the *ter* proximal DR. “o” and “p” indicate presence of a 5’-hydroxyl or 5’-phosphate, respectively, at the indicated end of the PCR fragment. (D) Constructs with the mutation next to the *ori* proximal side. “o” and “p” indicate presence of a 5’-hydroxyl or 5’-phosphate, respectively, at the indicated end of the PCR fragment. (E) Varying the distance between the mutation and the cassette (= size of the DR) or between the mutation and the end of the PCR fragment. Numbers below the different “modules” at the end of the PCR fragment indicate the size in bp of those “modules”. Red shaded areas indicate a strategy or combination that resulted in reduced success rate (compared to the other tests in the same series) while the green shaded area indicate a combination that resulted in improved success rate. Numbers above the DNA maps indicate the length (in bp) of the corresponding “module” in experiments where those sizes were kept constant.

#### The recombinogenic homology arms are not equal

When the strategy described above was used (Fig 6B and S1A Fig) a significant fraction of the transformants segregated to both the possible alleles, acting as heterozygotes with different alleles in the different DRs. The experiment was repeated with combinations of PCR fragments that would introduce the mutation in only one of the DRs to test if there was any difference in how often the mutation from the PCR product was introduced depending on at which side of the cassette it was located. The results of these transformations were that most transformants acted like heterozygotes, but a significant fraction was homozygous for the parental allele (Fig 6B). When the mutation was present on the homology arm closer to the origin of replication (*ori* proximal) 13% of the recombination events failed to introduce the mutation, while when it was present on the homology arm closer to the terminus of replication (*ter* proximal) 31% of the recombination events failed to introduce the mutation. Apparently, the mutation was lost more often on one homology arm than on the other during recombination. This may indicate a fundamental difference in the recombination events at the different ends. Based on these experiments it was not possible to determine if the bias in co-transfer of the mutation was dependent on the direction of replication (i.e. *ori* proximal versus *ter* proximal location of the mutation) or dependent on some other parameter such as transcription direction. To distinguish between these possibilities, both possible DIRex intermediates for introducing single point mutations into the genes *rpoS* and *clpA* were generated (S2 Fig). These genes have the same direction of transcription but are located on different replichores, which allows separation of any effects due to the direction of replication or transcription. While the efficiencies of successful co-transfer of the intended mutation varied between the two loci, in both cases the efficiency was greater when using a DIRex intermediate with the mutation in the *ori* proximal than in the *ter* proximal homology arm. Thus, in three genomic locations the efficiency of co-transfer of a mutation appeared to be influenced by the orientation of replication.

To examine this observation further and potentially find a more efficient method, an alternative strategy was tested (Figs 3C and 6C – E, and S1B Fig). Instead of trying to introduce the mutation at both sides of the cassette with the mutation in the middle of the flanking DRs, the DRs were chosen so that the mutation was left just next to the DR on only one side of the cassette. The effect of the different homology arms was tested by constructs with the mutation either on the *ori* proximal or the *ter* proximal PCR fragment. Consistent with the results from the first strategy, the transformations with the mutation on the *ori* proximal fragment (∼90%; Fig 6D – E) were more efficient at generating the desired mutation compared to the alternative (∼60%; Fig 6C).

#### The inequality of the homology arms can be overcome or aggravated by 5’-phosphorylation of the *ter* proximal fragment

Previous studies suggest that λ Red recombineering with dsDNA requires one of the strands of the incoming DNA to be completely degraded by Exo (Red α), leaving the other to act as substrate in the recombination [23,24]. To further characterize the unequal outcomes of recombination at the different ends, the strong preference of Exo for 5’-phosphorylated dsDNA ends [25] was used. If only one strand is phosphorylated that strand will be the preferred target for degradation, making the other strand more likely to have an intact 5’-end that can participate in the recombination. PCR products that were 5’-phosphorylated on either strand were generated and the segregation patterns of transformants were tested (Fig 6C and D). Phosphorylation of either end of the *ori* proximal PCR fragment had only minor effects on the success of introducing the mutation (Fig 6D). At the *ter* proximal end (Fig 6C), phosphorylation of the strand with 5’-homology to the lagging strand template had a strong detrimental effect on the success rate (reduced from 61 to 36%) while phosphorylation of the other strand increased the success rate (to 86%). Finally, the effect of the distance between the mutation and the cassette was tested by varying the size of the generated DRs, and the effect of the distance between the mutation and the end of the PCR fragment was tested by an alternative design, keeping the total size of the recombinogenic extension at 40 bp (including the DR, Fig 6E). In this test, the distance between the mutation and the cassette (i.e. the DR size) did not affect the outcome of the recombineering step, while the distance to the end of the PCR fragment did. When the mutation was 20 bp from the end of the fragment, the frequency was similar to when it was 40 bp from the end, but shorter distances led to reduced frequencies.

All in all, the preferred strategy for introducing a point mutation using DIRex is the one illustrated in Fig 3C. As the size of the DR affected the excision frequency (Fig 4B), but had no effect on the frequency of co-transfer of the mutation (Fig 6E), a long (25 bp or longer) DR is preferable. Due to the observed differences in recombination at the two ends it is preferable to have the mutation at the *ori* proximal side of the cassette. In other words, it is best to place the mutation in the PCR-fragment that has a 3’-end that is complementary to the lagging strand template, otherwise phosphorylation of one 5’-end may be needed to improve co-transfer (Fig 6C – D). Additionally, the mutation should be ∼20 or more bp away from the end of the PCR product. Using this strategy, 80% or more of the tested transformants contained the desired allele and could only segregate to this allele.

## Discussion

There are several strategies for using λ Red recombineering to generate marker-free and scar-free mutants. Methods using single-stranded DNA (ssDNA) oligonucleotides instead of double-stranded DNA (dsDNA) allow introduction of point mutations without the need for leaving any marker or scar sequence behind [8–10]. However, ssDNA recombineering suffer the drawback of not having any directly selectable phenotype. Unless the introduced mutations have selectable phenotypes on their own accord, they cannot easily be transferred between strains and hence further strain constructions become problematic. A compromise that works for non-essential genes is to combine dsDNA recombineering using a selectable and counter selectable marker with ssDNA recombineering [12–14]. In one method, a resistance cassette containing the recognition site for the homing endonuclease I-SceI is placed between small directly repeated copies of the target sequence, containing the desired mutation [11]. After introduction of the cassette it can be efficiently removed by co-expression of λ Red and I-SceI. This method requires curing the first λ Red plasmid and transformation with another plasmid that in addition to the λ Red genes carry the gene encoding I-SceI [6,11]. An alternative approach is to use CRISPR/Cas9 to make a cut in the wild-type allele after recombineering (using ssDNA or dsDNA), allowing only the designed mutant to survive [7]. CRISPR/Cas9 has a few shortcomings that makes DIRex competitive in many cases. Firstly, CRISPR/Cas9 requires the expression of a designed guide RNA (gRNA) that directs the cleavage of the wild-type allele. Synthesis and cloning of the gRNA genes has to be done specifically for each individual construct. Our method does not require any cloning step, and only requires two PCR reactions to generate the needed material. Secondly, Cas9 only cleaves within a certain distance from a protospacer adjacent motif (PAM). While PAMs are typically frequent it still means not all sites are equally accessible for cleavage by Cas9 [26].

Thus, the strategies described above either require (i) screening in the absence of a selectable phenotype, or require (ii) several transformation steps [6–14], or (iii) expression of additional factors to induce chromosome breaks (e.g. I-SceI or CRISPR/Cas9 plus a guide RNA; [7,12–14].

The strategy developed by Tear *et al.* [19], essentially adopting MIRAGE [18] for *E. coli*, removes the need for several transformation steps by using artificial gene-specific IRs (∼600 – 800 bp), located between gene-specific DRs. The large IRs enhance the rate of spontaneous and precise excision of the inserted cassette without the need for expression of any extrinsic recombinase. A selectable and counter selectable cassette allows for simple isolation of clones that have lost the insertion [19]. This method has the advantage that it can generate the desired mutation with only one transformation step, followed by selection for clones that have lost the transient insertion. The need to construct locus-specific IRs by OE-PCR for every locus is a drawback. The process could become unnecessarily cumbersome and time-consuming in cases where the OE-PCR needs to be specifically optimized for a particular locus. DIRex overcomes this limitation by not requiring any OE-PCR, which was enabled by our recent observation that recombineering with two overlapping “half-cassette” PCR products is as efficient as recombineering with the complete cassette as a single PCR product [20]. In addition, the fact that the IRs are locus derived limits the Tear *et al*. method [19] to only making deletions (since both copies of the IR sequence are by necessity lost in the process), and limits the minimum size of deletions to between ∼600 and 800 bp due to the size dependency of IR size and excision frequency. By using cassette derived IRs, DIRex does not suffer this limitation. Consequently, DIRex can be used for making deletions of any size, as well as replacements, exchanges and insertions, ranging in size from what can be practically included in a PCR oligo down to single nucleotides. For a comparison between DIRex and the *Tear et al*. method [19], see S4 Fig.

### Implications for the mechanism of λ Red mediated dsDNA recombineering

Red mediated recombination between relatively short linear dsDNA and chromosomal targets is biased towards the lagging strand, suggesting that the recombination is initiated by annealing of a 3’-end of the incoming DNA to a single-stranded gap between Okazaki fragments during lagging strand synthesis [23,24]. Mosberg *et al.* [24] used dsDNA cassettes with strand-specific mismatches to examine the mechanism of Red mediated dsDNA recombination. They found that the majority (∼72%) of the recombinants inherited sequences from both ends of the strand which was complementary to the lagging template strand, while a minority of the recombinants contained sequences only from the other strand or from both strands at about equal frequencies. These results were taken as support of a fully single-stranded intermediate in recombineering, and the minor type of recombinants were interpreted as a second recombination event after the successful recombination with the lagging targeting strand. However, the distribution of their recombinant types (68:7:9:10) is also consistent with two competing mechanisms of recombination. A lagging strand biased mechanism resulting in most of the major type of recombinants as suggested, and another mechanism apparently lacking any strand bias accounting for an approximately equal distribution of all four types of recombinants. Another observation that points towards two mechanisms is that strand bias is gradually lost when the size of the incoming DNA is increased from ∼1 to ∼3 kb, and concurrently transformation frequencies drop to a plateau [23]. Recombineering with two overlapping PCR fragments require at least two DNA strands, that need to be nearly intact in both ends to have enough complementarity both to each other and to their chromosomal target sequences. This implies that both the leading and lagging strands need to be involved in the recombination, which would remove or obscure any strand bias. Our recent observation that overlapping fragments recombine with similar frequencies as single fragments of the same total size (3.3 kb; 18) may indicate that even when transforming with a single PCR product, a sizeable proportion of the incoming DNA is fragmented into smaller pieces before recombination, and that recombination needs to occur between more than one fragment to generate a viable transformant. Observations made during the development of DIRex may provide clues to this alternative mechanism of Red recombination. Firstly, a difference in success rates was seen depending on which side of the cassette the mutation was located, and this difference appeared to be linked to the orientation relative to the direction of replication (Fig 6 and S2 Fig). Secondly, the success rate when the mutation was on the *ter* proximal side of the cassette could be slightly improved by 5’-phosphorylation of the strand that is complementary to the leading template strand, but severely reduced when the strand complementary to the lagging template strand was 5’-phosphorylated. When the mutation instead was on the *ori* proximal side of the cassette phosphorylation of either strand of the mutation-containing fragment caused a relatively small reduction in the success rate (Fig 6C). Our observations are consistent with a strand unbiased recombination mechanism. If 5’-phosphorylation of one of the four DNA strands results in complete degradation of that strand, its complementary strand would not be substrate for degradation by Exo and consequently be used more frequently in the recombination. The poor frequency of co-transfer when only the strand with 3’-complementarity to the leading strand template is available to recombine on the *ter* proximal side of the insertion could give a hint towards different homology requirements at the different ends or the timing of recombination events. Perhaps recombination involving a 3’-end that is annealed to a gap in the leading strand does not always use the full 3’-end. Alternatively, the recombination events occur in a set order, leaving that 3’-end exposed to an endogenous 3’→5’ exonuclease activity for a longer time, allowing it to be trimmed to the point where the mutation is not co-transferred.

With DIRex a high rate of successful mutant isolation is achieved, with a single transformation and minimal cost and effort in terms of construction, screening and sequencing. It is demonstrated that DIRex is useful for generating deletions, for producing single nucleotide substitutions and for replacing shorter or longer sequences with short synthetic sequences. Although not demonstrated here, this makes DIRex useful for introducing affinity tags, degradation tags, or other short sequences as long as they are short enough to be included in a DNA oligo. Additionally, the method should be adaptable to use with BAC clones, or in other bacterial species, as long as the necessary genetic tools (λ Red or similar recombineering technology, and counter-selectable cassettes) are available.

## Materials and Methods

### Strains and growth conditions

All *Salmonella* strains are derivatives of *S. enterica* serovar Typhimurium strain LT2 and all *Escherichia coli* strains are derivatives of *E. coli* K12 strain MG1655. All strains are listed in S1 Table. Generalized transduction using phages P22 HT 105/1 *int-201* [27] and P1 *vir* [28] were used to move chromosomal markers between strains. For rich medium either SOC (20 g/L tryptone [oxoid], 5 g/L yeast extract [oxoid], 0.5 g/L NaCl, 0.25 mM KCl, 10 mM MgCl_2_, 4 g/L glucose) or LB (10 g/L NaCl, 10 g/L tryptone [oxoid] and 5 g/L yeast extract [oxoid]) was used. The LB was supplemented with 15 g/L agar (oxoid) to make LB agar (LA) plates. Sucrose-selection plates (LA without NaCl, supplemented with 50 g/L sucrose) were used to select against cells expressing the *sacB* gene. M9 minimal media [29] with 0.2% (w/v) glucose was supplemented with L-histidine (0.1 mM; Sigma-Aldrich) or L-tryptophan (0.1 mM; Sigma-Aldrich) when appropriate. Antibiotics (Sigma-Aldrich) were used at the following concentrations: chloramphenicol (cam), 6.25 mg/L; tetracycline (tet), 7.5 mg/L.

### λ Red recombineering

PCR reactions to generate DNA for recombineering were performed with Phusion DNA polymerase (Thermo Fisher), using a PCR program described previously [20]. All oligo sequences are listed in S2 Table. For all constructions, primers with names ending with “-fP1” were used together with the primer *cat*-midF and the template *Acatsac3* (GenBank: MF124799), and primers ending with “-rP1” were used together with *cat*-midR2 and the template *Acatsac1* (GenBank: MF124798). For making cells competent for λ Red transformation through electroporation, cultures of strains containing the pSIM5-Tet plasmid [30] were grown overnight at 30°C in salt free LB (LB without NaCl) with 2 g/L glucose and 7.5 mg/L tetracycline. Cultures were diluted 1:100 in the same medium, pre-warmed to 30°C, and grown at 30°C until OD_600_ ≈ 0.2 (measured with 1 cm light path in a Shimadzu UV mini 1240 spectrophotometer). Once the target OD was reached, the culture flasks were moved to a 42°C shaking water bath to induce expression of the temperature-controlled λ Red genes. After 15 min at 42°C (OD_600_ ≈ 0.3) the cultures were cooled in an ice-water bath for at least 10 min. The cells were pelleted by centrifugation at 4°C (4,000×g, 7 min) and all the medium was removed. The cells were washed once in ice-cold 10% glycerol (1/4 of the culture volume), pelleted (4,000×g, 6 min) and re-suspended in ice-cold 10% glycerol (320 μl per 25 ml initial culture volume). Electro-competent cells (20 μl) and DNA (about 0.15 pmol PCR product) were mixed in electroporation cuvettes (1 mm gap, Bio-Rad) on ice, and electroporated in a Gene Pulser Xcell (Bio-Rad) at 2.5 kV, 400 Ω and 25 μF. After electroporation, cells were immediately moved to pre-warmed (42°C) SOC in a 42°C water bath and incubated without shaking for 15 min before plating on LA plates with 12.5 mg/L chloramphenicol. The short recovery was used to avoid growth and prevent the formation of siblings prior to plating, enabling accurate determination of the ratios of different classes of transformants. In cases when siblings were not considered a problem the cells were allowed to recover at 37°C for several hours or overnight.

### Phosphorylation of 5’-ends

To generate PCR products with one phosphorylated 5’-end, one of the oligonucleotides for the PCR reactions were phosphorylated using T4 polynucleotide kinase (PNK, New England Biolabs). Briefly, 100 pmol oligo was incubated for 30 min at 37°C with 20 nmol ATP and 10 units PNK in 20 μl 1x reaction buffer A for PNK. After heat inactivation of the enzyme (20 min at 80°C) the oligo was used as primer in PCR in combination with a non-phosphorylated primer.

### Curing strains from DIRex intermediates

One single (blue, Cam^R^) colony per isolated clone from a transformation or transduction was streaked out for single colonies on sucrose selection plates. After incubation over night at 37°C this resulted in growth of white, Suc^R^, Cam^S^ colonies, sometimes mixed with a few blue colonies. Blue, Suc^S^ clones still contained at least part of the *cat*-*sacB* cassette and were discarded as false positives. White, Suc^R^ clones were confirmed by PCR and sequencing. A small minority (typically <1%) of transformants displayed either stronger or weaker blue color, indicating a different expression level of amilCP. Some of these clones segregated to Suc^R^ while others did not (data not shown), indicating they were erroneous recombinants or contained mutations within the *AcatsacA* cassette. These recombinants were excluded from analysis by discarding clones that differed visibly in color.

### Segregation frequency assay

Cultures were started from six independent colonies of each transformant clone in LB supplemented with chloramphenicol. After overnight growth at 37°C the cultures were diluted 1:5×10^6^ into the same medium (to a final culture density of approximately 10^3^ cells/ml) and allowed to grow overnight again. This population bottleneck was done to minimize the risk of preexisting mutants in the cultures. Serial dilutions of these cultures were plated on sucrose selection plates to determine the number of sucrose resistant segregants (white) and false positives (blue), and on LA plates to determine the culture densities (which varied between 2.9 – 6.1×10^9^ cfu/ml with an average of 4.9×10^9^ cfu/ml).

## Acknowledgements

This study was funded by grants from the Swedish Research Foundation (VR). I am very grateful to Anna Knöppel, Hervé Nicoloff, Dan I. Andersson and Erik Gullberg for critical reading and helpful comments on this manuscript.

## Supporting information

**S1 Fig.**
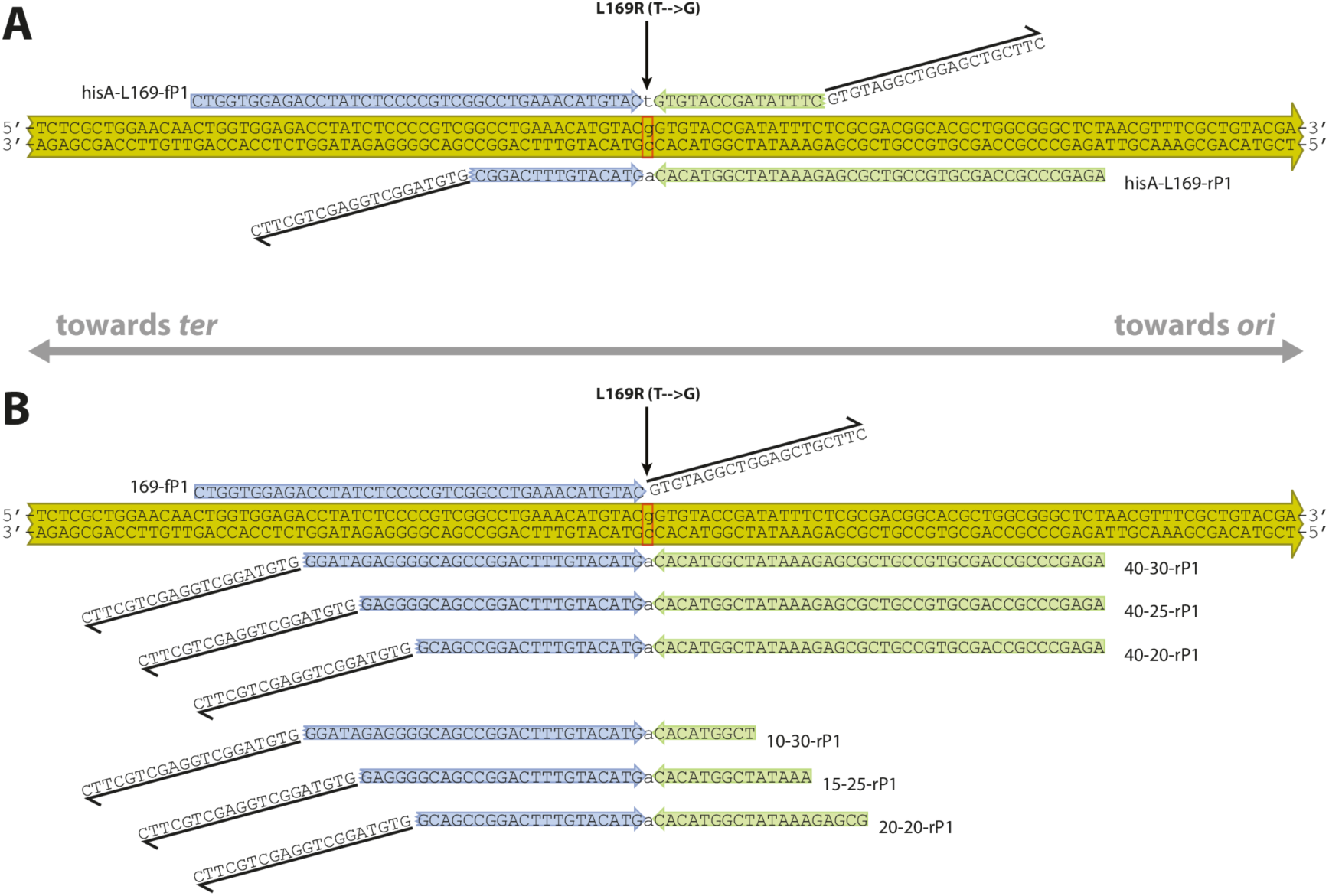
Examples of primers used for DIRex. Oligos are aligned to their homologous sequences in the *Salmonella enterica hisA* gene. Part of the coding sequence of *hisA* is shown as a double-stranded sequence on top of a yellow arrow, showing the direction of the gene. The vertical arrow points towards the position of the L169R (CAG to CGG) mutation, which is also highlighted with a red box and lower case letters. The green and blue arrows correspond to the “modules” of the recombinogenic ends that are highlighted in green and blue in Fig 6. The slanted “half-arrow” sequences indicate the template annealing portions, which are identical in all oligos. (A) Primers used for the constructs used for the experiments described in Fig 4 and 6B. The overlap between the “upper” primer (hisA-L169-fP1) and the “lower” primer (hisA-L169-rP1) generates the DR in the transformants. (B) Alternative primers used for the constructions in Fig 6D and E. The upper primer (169-fP1) was used with one of the lower primers (NN-MM-rP1) to generate different sized DRs from the sequences that overlap between the lower and upper primer, and different amounts of homology to the “left” of the mutation. The design of the oligos used for the constructs in Fig 6C is similar to 40-25-rP1 and 169-fP1, but target the opposite strands to place the cassette on the other side of the mutation.

**S2 Fig.**
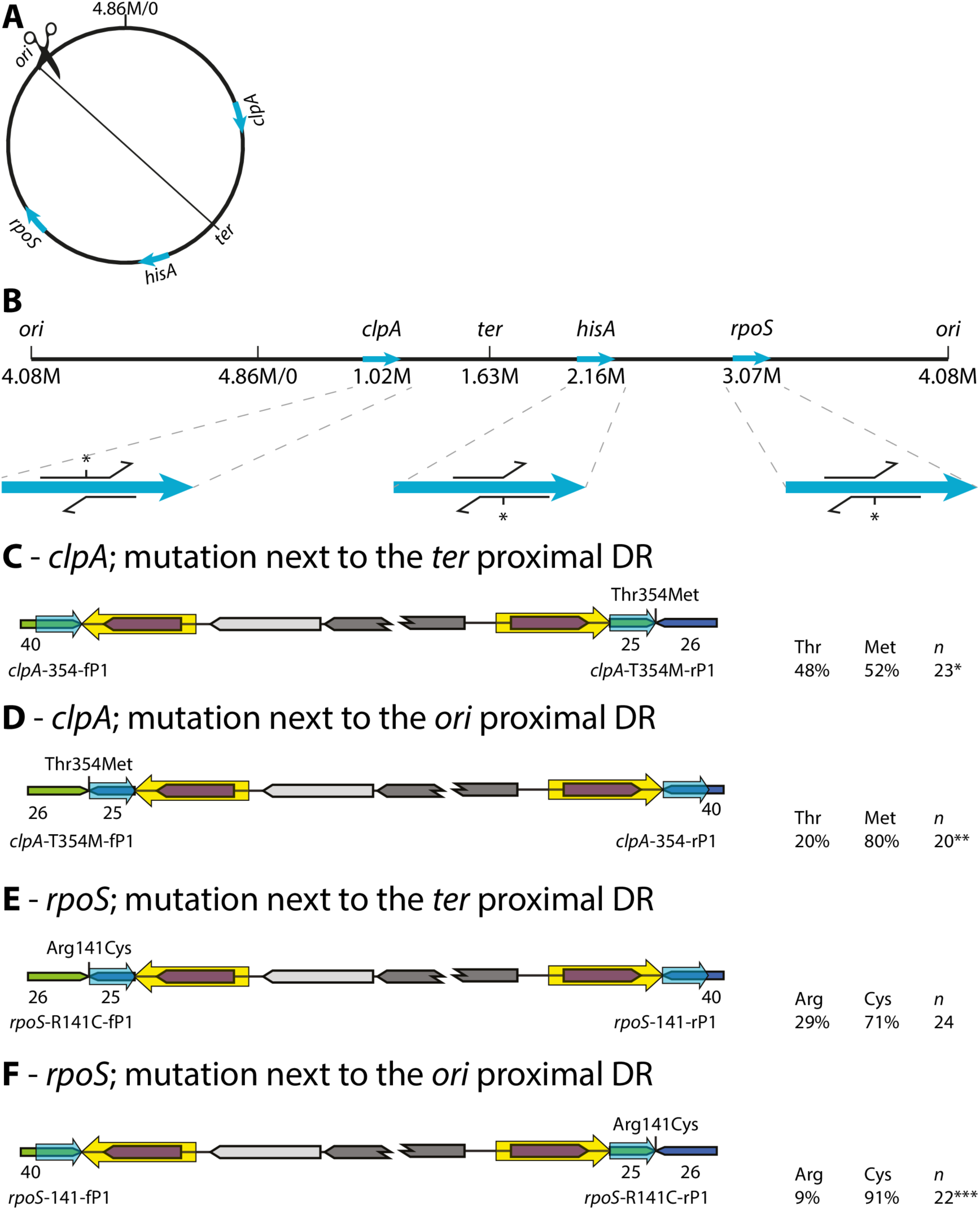
The efficiency of generating point mutations depends on the direction of replication. (A) Circular map of the *S. enterica* chromosome with the directions and approximate locations of the *ori* and *ter* and the three genes *clpA*, *hisA* and *rpoS*. (B) Linear representation of the *S. enterica* chromosome, starting and ending at *ori*. Zoomed in regions show the individual *clpA*, *hisA* and *rpoS* genes as light blue arrows. The most efficient locus specific primers are indicated, with the mutation-containing primer indicated with an asterisk (these oligos were used in S2D and F Fig, and Fig 6D and E. Note that in all three examples the mutation is on the oligo whose homology region is directed towards the *ori* proximal side. (C) Transformation to construct a Thr354Met mutation in *clpA*, using a DIRex intermediate with the mutation in the *ter* proximal DR. (*) Out of 24 sequenced Suc^S^, white segregant clones, one had a ∼400 bp deletion in *clpA* and was discarded. (D) Transformation to construct the same mutation as in (C), but using a DIRex intermediate with the mutation in the *ori* proximal DR. (**) Out of 24 sequenced Suc^S^, white segregant clones, one had a frameshift mutation in a neighboring codon and was discarded. Three produced poor sequence but was not further tested. (E) Transformation to construct an Arg141Cys mutation in *rpoS*, using a DIRex intermediate with the mutation in the *ter* proximal DR. (F) Transformation to construct the same mutation as in (E), but using a DIRex intermediate with the mutation in the *ori* proximal DR. (***) Out of 24 sequenced Suc^S^, white segregant clones, two had frameshift mutations in neighboring codons and were discarded. The direction of transcription of both genes is left to right in the picture.

**S3 Fig.**
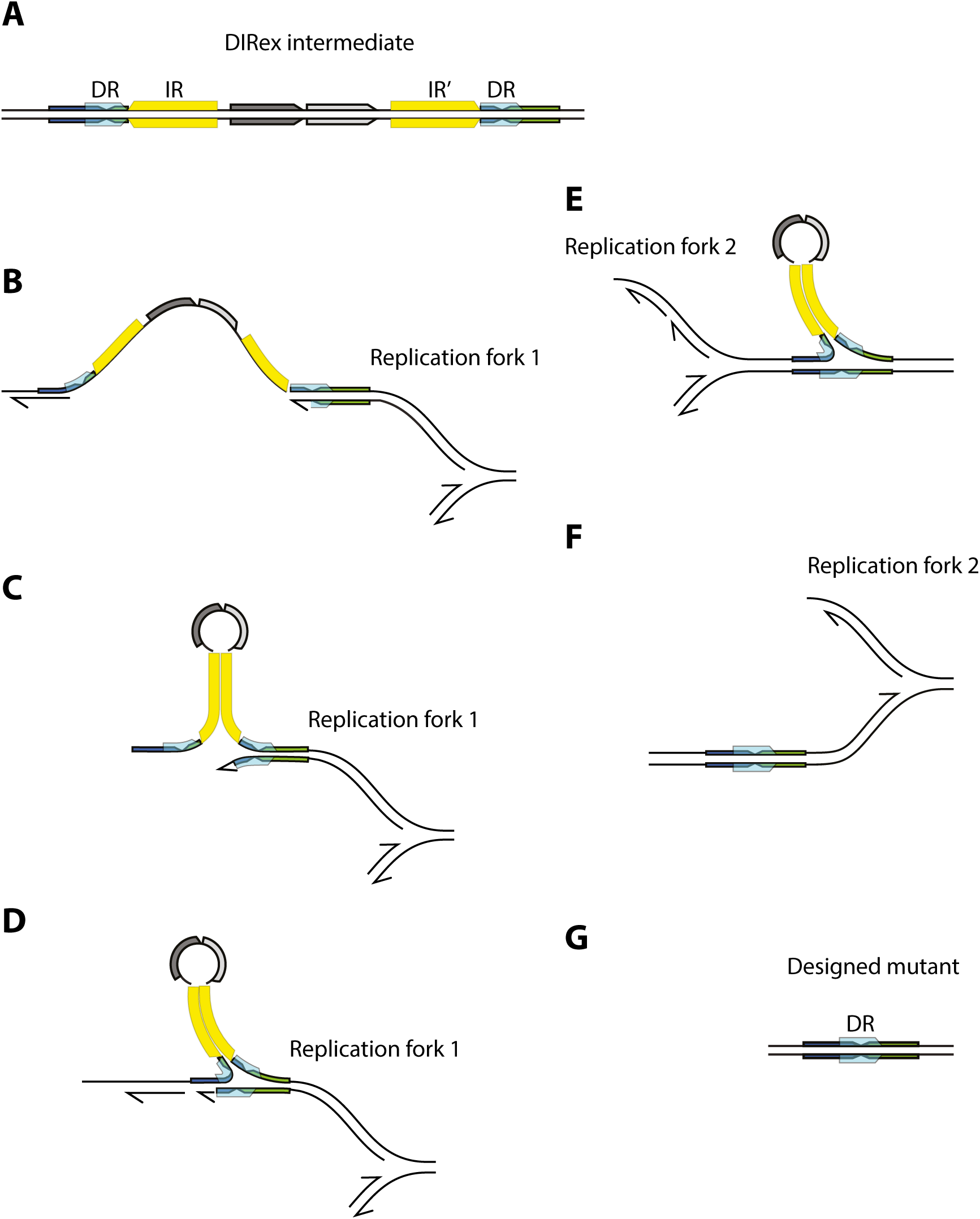
Excision of a DIRex intermediate through slipped-strand misparing during lagging strand replication. (A) A DIRex intermediate. DRs are indicated as turquoise boxes and IRs as yellow boxes. IR’ indicates the reverse complement of IR. (B) Complementary sequences from IR and IR’ may be exposed during passage of a replication fork, depicted here as a long continuous gap between Okazaki fragments on the lagging strand. (C) Transient intra-strand basepairing between IR and IR’ forms a large stem-loop, stalling replication and bringing a DR and its complementary sequence next to each other. (D) Pairing of the DR and its complementary sequence allows continued lagging strand synthesis. (E) After passage of the first replication fork the new leading strand template lacks the DIRex intermediate and has only one copy of the DR sequence. (F) After passage of the next replication fork, the replicated leading strand carries the designed mutation. (G) The designed mutation segregates into one of the daughter cells during cell division. The model is essentially as suggested by Bzymek and Lovett [31] for stimulation of deletion by uninterrupted palindromes through misalignment during lagging strand synthesis.

**S4 Fig.**
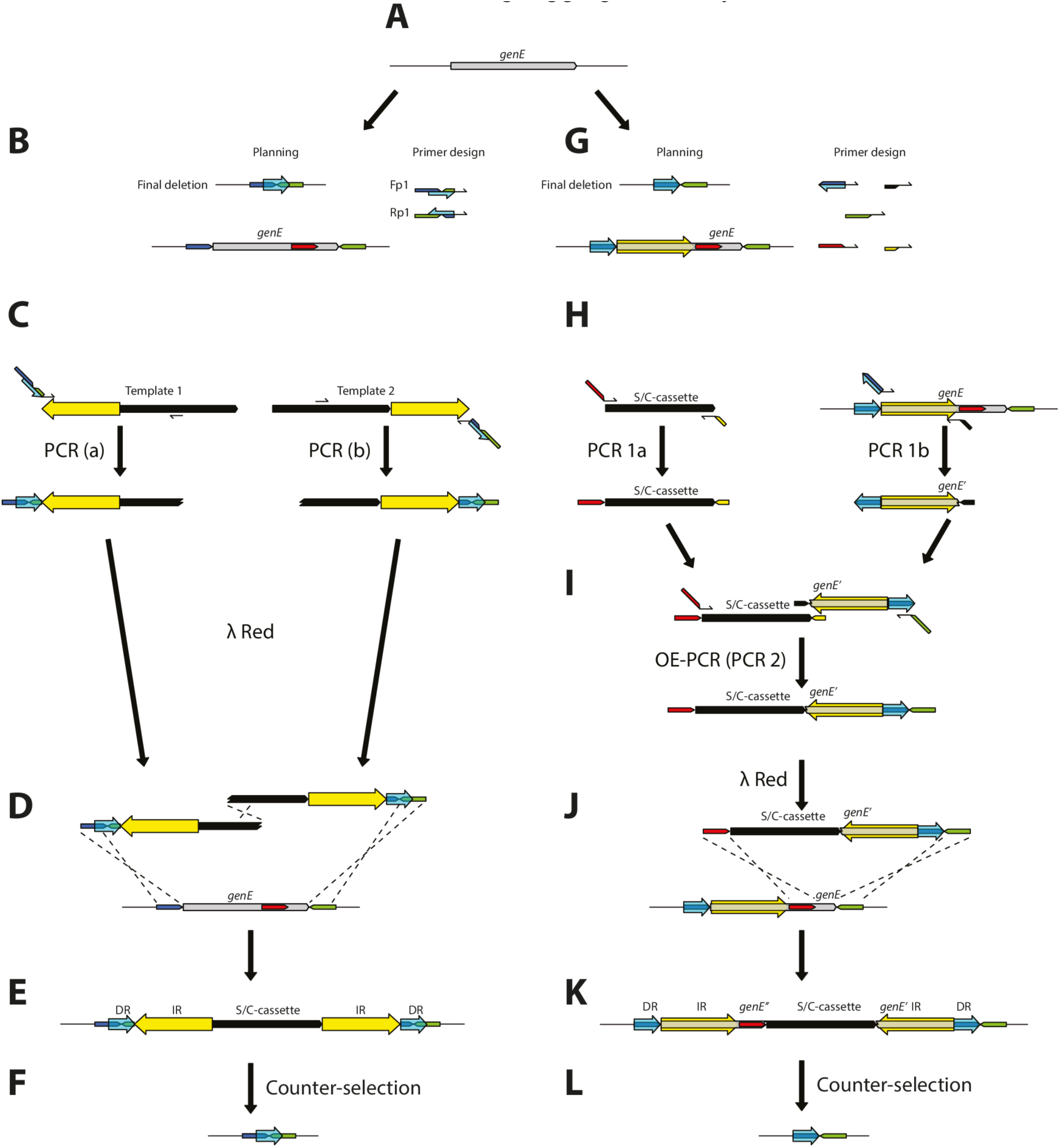
Comparison between DIRex and a similar method described by Tear *et al*. Both methods are used for the construction of the same precise deletion of the hypothetical gene “*genE*”. (B – F) Use of DIRex for deletion of a gene. (B) Only two gene-specific oligos are needed, in combination with two cassette-specific primers. Recombinogenic tails of the locus-specific primers are chosen so that both oligos contain the designed deletion junction, in this case bringing the “blue box” sequence next to the “green box” sequence and deleting the entire “*genE*” sequence. The DRs will thus consist of part of the “blue box” and part of the “green box”. (C) Two separate PCR reactions are used to amplify two overlapping “half-cassettes”, each containing one copy of the ∼800 bp IR sequence, as well as complementing portions of a selectable and counter selectable (S/C) cassette. (D) The two PCR products are mixed in equimolar amounts and transformed into λ Red induced cells. (E) A semi-stable DIRex intermediate is formed. The S/C cassette (black) is flanked by two inverted repeat sequences (yellow, IR), which in turn are flanked by directly repeated sequences (turquoise, DR), each containing the deletion junction. (F) The final deletion mutant is isolated through selection against the S/C cassette. (G - L) The method described by Tear *et al*. [19] for deleting a gene. (G) Five specifically designed primers are needed. In addition to generating locus-specific DRs, locus-specific IRs has to be chosen, which limits the method into only constructing deletions and limits the minimal deletion size to the size of the IRs. (H) Two PCR reactions generates the S/C cassette and an artificial locus derived IR-DR cassette. (I) The overlapping S/C-cassette and IR-DR cassette is joined through overlap extension PCR (OE-PCR). (J) The OE-PCR product is transformed into λ Red induced cells. (K) A semi-stable intermediate, analogous to the DIRex intermediate in (E) is formed. (L) The final deletion mutant is isolated through selection against the S/C cassette. Note that the end results in (F) and (L) can be identical even if the DRs, IRs and homology extensions of the primers used are different.

**S1 Table. Bacterial strains.**

**S2 Table. Oligonucleotides.**

